# Noncanonical inflammasome assembly requires caspase-11 catalytic activity and intra-molecular autoprocessing

**DOI:** 10.1101/2022.09.21.508824

**Authors:** Daniel C. Akuma, Daniel Grubaugh, Chukwuma E. Odunze, Sunny Shin, Cornelius Y. Taabazuing, Igor E. Brodsky

## Abstract

Inflammatory caspases are cysteine protease zymogens whose activation following infection or cellular damage occurs within supramolecular organizing centers (SMOCs) known as inflammsasomes. Inflammasomes are large oligomeric complexes that recruit caspases to undergo proximity-induced autoprocessing, leading to formation of the enzymatically active form that cleaves downstream targets. Binding of bacterial LPS to its cytosolic sensor, caspase-11 (Casp11), promotes Casp11 aggregation within a high molecular weight complex known as the noncanonical inflammasome, where it is activated to cleave gasdermin D and induce pyroptosis. However, the cellular correlates of Casp11 oligomerization and whether Casp11 forms an LPS-induced SMOC within cells remain unknown. Using fluorescently labeled Casp11, we found that LPS transfection of macrophages induced Casp11 speck formation, providing direct evidence that Casp11 forms LPS-induced specks in macrophages. Unexpectedly, we found that catalytic activity was required for Casp11 to form LPS-induced specks in macrophages. Importantly, both catalytic activity and autoprocessing were required for Casp11 speck formation in an ectopic expression system, and Casp11 processing via an exogenous protease was sufficient to induce Casp11 speck formation. These data reveal a previously-undescribed role for Casp11 catalytic activity and self-cleavage in non-canonical inflammasome assembly and indicate that Casp11 catalytic activity and autoprocessing occur upstream of, and mediate, Casp11 oligomerization. Our data provide new insight into the molecular requirements for assembly of Casp11 noncanonical inflammasome complexes.

## INTRODUCTION

The mammalian innate immune system relies on evolutionarily conserved pattern recognition receptors (PRRs) to detect microbe-derived pathogen-associated molecular patterns (PAMPs) and rapidly respond to potential microbial threats (Blander and Sander, 2012; Janeway, 1989; Janeway and Medzhitov, 2002). Such rapid responses are critical for efficient host defense in the face of pathogens that have the capacity to replicate rapidly and suppress or evade immune detection. A common feature of PRR activation is their inducible assembly into higher order oligomeric protein complexes, termed supra-molecular organizing centers (SMOCs), which mediate the effector function of the PRR signaling pathway in response to microbial infection or pathologic stimulus (Kagan et al., 2014).

Caspases-1 and −11 are inflammatory caspases that are recruited into SMOCs known as inflammasomes in response to the presence of pathogen-derived signals or molecules (Broz and Dixit, 2016; Lamkanfi and Dixit, 2014). Inflammatory caspases are zymogens that contain a C-terminal enzymatic domain and an N-terminal caspase activation and recruitment domain (CARD), which promotes SMOC formation through homotypic interactions with other CARD-containing adapter proteins as well as homodimerization with each other (Lu et al., 2014; Park et al., 2007; Ting and Davis, 2005). The C-terminal enzymatic domain of caspases is comprised of large (17-20 kDa) and small (10-12 kDa) subunits separated by an inter-domain linker (IDL), which contains the self-cleavage site that undergoes autoprocessing following caspase oligomerization (Ross et al., 2022; Walker et al., 1994; Wilson et al., 1994).

Caspase-1 is recruited to complexes known as canonical inflammasomes that contain a sensor protein of the NLR family, the adaptor protein ASC, and caspase-1 itself (Broz and Dixit, 2016; Lamkanfi and Dixit, 2014). In contrast, caspase-11 (Casp11) binds directly to lipopolysaccharide (LPS) from gram-negative bacteria (Shi et al., 2014), and does not require a known sensor or adapter for activation of pyroptosis via a pathway termed the non-canonical inflammasome (Kayagaki et al., 2011). Recruitment of Casp1 and Casp11 to their respective SMOCs results in their autoprocessing, leading to activation of their ability to cleave downstream targets via a process termed proximity-induced activation (Salvesen and Dixit, 1999; Shi, 2004). Casp11-dependent cleavage of its target, gasdermin D, leads to lysis of the cell, as well as release of inflammatory contents, triggering pyroptosis, a pro-inflammatory programmed cell death (Aglietti et al., 2016; Kayagaki et al., 2015; Shi et al., 2015). As a result, Casp1 and Casp11 play important roles in antimicrobial host defense, but can also promote pathologic autoinflammatory disease when dysregulated (Franchi et al., 2009; Hagar et al., 2013; Henao-Mejia et al., 2012; Kayagaki et al., 2013).

Inducible dimerization is used to model assembly of functional caspase complexes within cells (Ball et al., 2020; Boucher et al., 2018; Oberst et al., 2010; Ross et al., 2018). Caspases whose N-terminal domain is replaced with a dimerizable domain, such as FKBP506, undergo autoprocessing and exhibit catalytic activity toward their substrates in the presence of dimerizing agents. Interestingly, Casp11 mutants whose autoprocessing site is ablated, have significantly reduced protease activity toward their substrates and have defective ability to induce cell death, despite being dimerized and having an intact catalytic site (Ross *et al*., 2018). Furthermore, CRISPR targeting of Casp11 autoprocessing or catalytic activity demonstrates that both are important for Casp11-dependent pyroptosis and lethal sepsis (Lee et al., 2018).

The observation that autoprocessing is needed to generate a fully functional Casp11 complex despite being dimerized and catalytically active (Ross *et al*., 2018) implies that formation of a functional Casp11 SMOC involves additional steps beyond inducible dimerization. Interestingly, while noncleavable Casp8 is active *in vitro* (Chang et al., 2003) or in the presence of kosmotropic salts (Pop et al., 2007), autoprocessing is also required for full Casp8 activity toward its substrates within cells, implying that autoprocessing stabilizes the active form of the enzyme (Oberst *et al*., 2010). This parallels recent findings with Casp1, which requires autoprocessing for cleavage of IL-1β (Broz et al., 2010) and GSDMD (Ball *et al*., 2020). However, while ASC fluorescent reporters have enabled the dynamic tracking of canonical inflammasome assembly within cells (Stutz et al., 2013; Tzeng et al., 2016), the cellular correlates of Casp11 oligomerization within cells in response to cytosolic LPS remain poorly defined.

Here, we find using fluorescent Casp11 reporter fusions that cytosolic LPS induced Casp11 assembly into large perinuclear specks, the first time to our knowledge, that LPS-induced noncanonical inflammasome speck assembly has been directly observed *in vivo*. Unexpectedly, Casp11 catalytic activity was required for LPS-inducible Casp11 speck formation in macrophages, suggesting that catalytic activity acts *upstream* of Casp11 oligomerization to facilitate assembly of the fully activated Casp11 SMOC. Intriguingly, both Casp11 catalytic activity and autoprocessing were required for spontaneous speck assembly in HEK293T cells, consistent with a model whereby higher order oligomerization of Casp11 requires catalytic activity and autoprocessing. Consistently, inducible IDL processing of Casp11 by an exogenous protease was sufficient to mediate speck assembly. Moreover, we find that intramolecular (*cis*) processing, rather than intermolecular (*trans*) processing of casp11 mediates speck formation. Altogether, we demonstrate that Casp11 catalytic activity and autoprocessing act upstream of Casp11 oligomerization to mediate assembly of the fully functional Casp11 SMOC. Our findings uncover a previously undescribed property of caspase-containing SMOCs and imply that Casp11 activation involves a feed-forward mechanism whereby self-processing of Casp11 facilitates assembly of the pyroptosis-competent Casp11 complex.

## RESULTS

### Caspase-11 catalytic activity is required for cytosolic LPS-induced speck formation

Upon sensing cytosolic LPS, Casp11 oligomerizes into high molecular weight complexes known as noncanonical inflammasomes (Shi *et al*., 2014). To better understand the oligomerization dynamics of the Casp11 inflammasome, we generated a lentiviral construct encoding C-terminal fusions of mCherry with full-length Casp11 (Casp11^WT^-mCherry) and the corresponding C254A mutant lacking catalytic activity, and transduced these constructs into *Casp11*^-/-^ bone marrow-derived macrophages (BMDMs) (**Fig. 1A**). Casp11^WT^-mCherry but not Casp^C254A^-mCherry restored dose-dependent GSDMD cleavage and cytotoxicity in *Casp11*^-/-^ BMDMs, indicating that the tagged mCherry construct retains function (**Fig. 1B**, **C**). We next observed Casp11 localization using confocal microscopy. As expected, both Casp11^WT^-mCherry BMDMs and Casp11^C254A^-mCherry-transduced cells exhibited diffuse cytoplasmic staining in the absence of LPS stimulation (**Fig. 1D**). Importantly, Casp11^WT^-mCherry coalesced into large mCherry specks in response to transfection of LPS, consistent with the concept that Casp11 assembles into an LPS-induced higher order complex, or SMOC (**Fig. 1D**). As expected, Casp11^WT^-mCherry exhibited dose-dependent speck formation in response to increasing intracellular LPS levels, paralleling the observations of dose-dependent toxicity (**Fig. 1E**). Surprisingly, Casp11^C254A^-mCherry did not form specks in response to LPS, even at higher doses, despite being expressed at similar or slightly higher levels (**Fig. 1C-E**). These findings imply that CARD-dependent binding of Casp11 to LPS is not sufficient to mediate Casp11 speck formation within cells, and that Casp11 catalytic activity facilitates assembly of the higher order noncanonical inflammasome.

**Figure 1.**
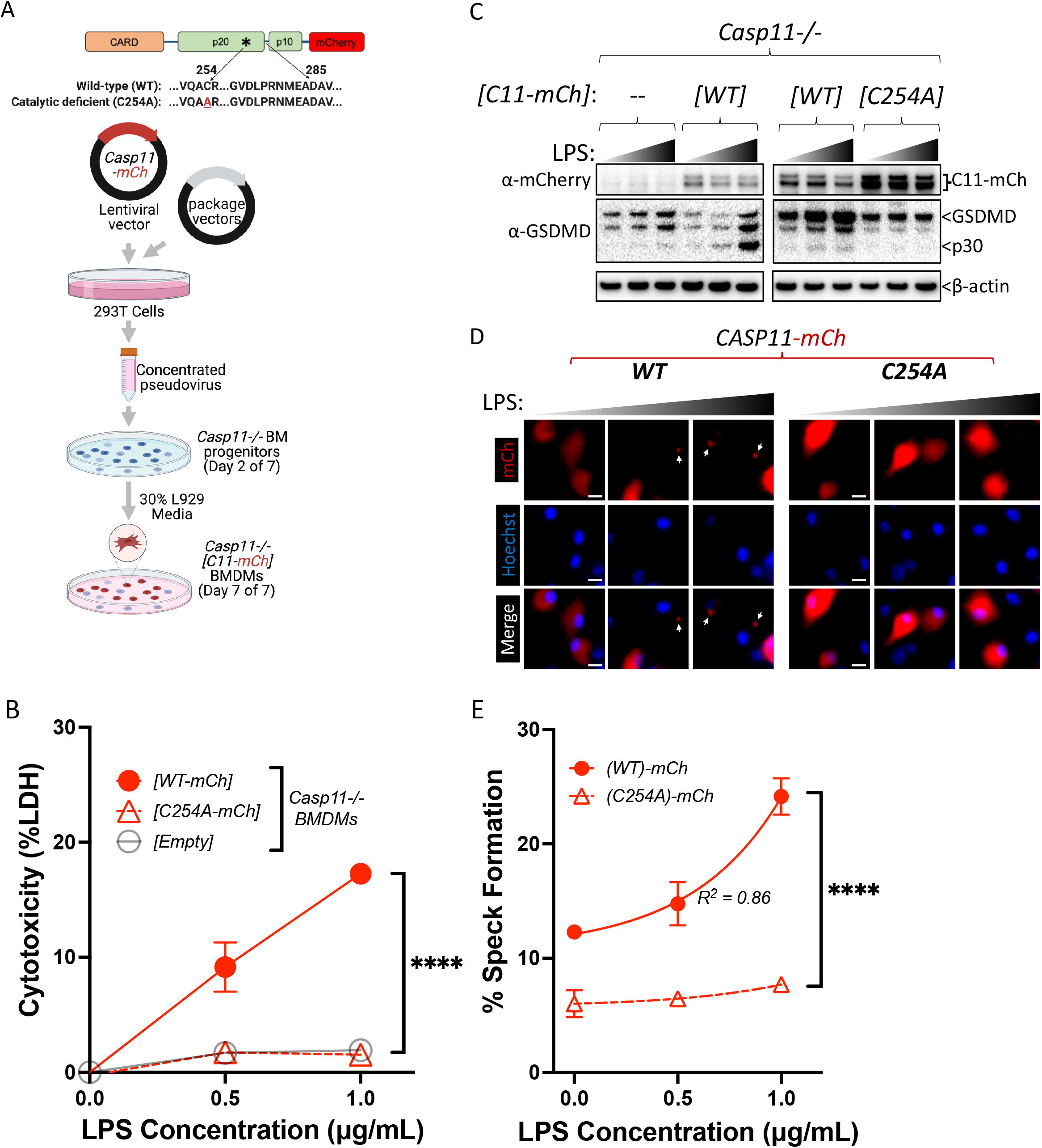
Caspase-11 catalytic activity is required for cytosolic LPS-induced speck formation. (A) Schematic describing complementation of *Casp11*^-/-^ BMDMs with mCherry-tagged caspase-11: Wild-type (*WT-mCh*) or Catalytic-deficient (*C254A-mCh*). (B-E) *Casp11*^-/-^ primary BMDMs with the indicated transgenes or empty-vector control were primed with Pam3CSK4 for 4 h, followed by transfection with LPS from *S. enterica* serovar Minnesota. (B) caspase11-mediated pyroptosis was assayed 16 hr later by LDH release (C) Casp11-mCherry expression and gasdermin D processing in response to LPS transfection was assessed by western blotting for mCherry and GSDMD as indicated. β-actin was used as a loading control. (D) Cells expressing indicated Casp11-mCherry constructs were fixed 6 hours post-LPS transfection and prepared for microscopy. Nuclei are stained with Hoechst. White arrows indicate Casp11-mCherry specks. Scale bar, 10 μm. (E) Speck formation from cells in D was quantified as percentage of mCherry-expressing cells containing a speck. Dose-response curves were plotted using non-linear regression (Log_2_; three parameters). Each data point represents 4 image frames (100-150 cells) per well and 3 wells per condition for a total of 300-450 cells. All error bars represent mean ± SEM of triplicate wells; representative of 2-3 independent experiments. *p <0.05, **p < 0.01, ***p < 0.001, ****p < 0.0001, ns not significant. Two-way ANOVA with Sidak’s multiple comparison test.

### Caspase-11 catalytic activity is required for spontaneous caspase-11 oligomerization in HEK293T cells

Our observation that catalytic activity was essential for LPS-induced aggregation in macrophages suggested that Casp11 catalytic activity somehow contributes to higher order complex assembly, rather than higher order complex formation being due entirely to Casp11 binding to LPS. To further dissect these mechanisms, we employed an extensively employed system to define the molecular basis for inflammasome assembly in HEK293T cells (Kayagaki *et al*., 2015; Rauch et al., 2017; Ross *et al*., 2018; Shi *et al*., 2014; Tenthorey et al., 2014). In 293T cells, ectopic Casp11 expression mediates activation of the non-canonical inflammasome and processing of GSDMD (Aglietti *et al*., 2016; Kayagaki *et al*., 2015; Shi *et al*., 2015). Consistent with previous observations, Casp11-mCherry was robustly expressed, and co-expression of Casp11^WT^-mCherry and GSDMD led to dose-dependent GSDMD cleavage and cell death that required Casp11 catalytic activity (**Extended Data Fig. 1**). In this system, Casp11^WT^-mCherry spontaneously assembled into large aggregates reminiscent of ASC-containing specks (Stutz *et al*., 2013; Tzeng *et al*., 2016), whereas Casp11^C254A^-mCherry remained diffuse within the cytosol, consistent with our previous finding in macrophages that catalytic activity of Casp11 is necessary to initiate or propagate Casp11 inflammasome assembly (**Fig. 2A, B**). Moreover, the pan-caspase inhibitor Z-VAD-FMK (zVAD) blocked spontaneous speck assembly and autoprocessing by Casp11^WT^-mCherry in a dose-dependent manner (**Fig. 2C-E**). Consistently, Casp11^C254A^-mCherry did not aggregate into visible specks, and its cytoplasmic distribution was not detectably altered in the presence of zVAD. Similar doses of zVAD abrogated LPS-induced Casp11-mediated pyroptosis in BMDMs (**Extended data Fig. 2A-C**). Importantly, consistent with prior studies (Shi *et al*., 2014), catalytic activity was not required for Casp11 self-association per se, as Casp11^WT^ and Casp11^C254A^ associate both with themselves and each other in reciprocal co-IP studies (**Extended Data Fig. 3**). Speck formation also occurred with a Casp11-citrine fusion, but not with a CARD only-citrine fusion, providing further evidence that the enzymatic domain is important in Casp11 oligomerization (**Extended Data, Fig. 4**). Together, these data imply that catalytic activity contributes to higher order Casp11 oligomerization.

**Figure 2.**
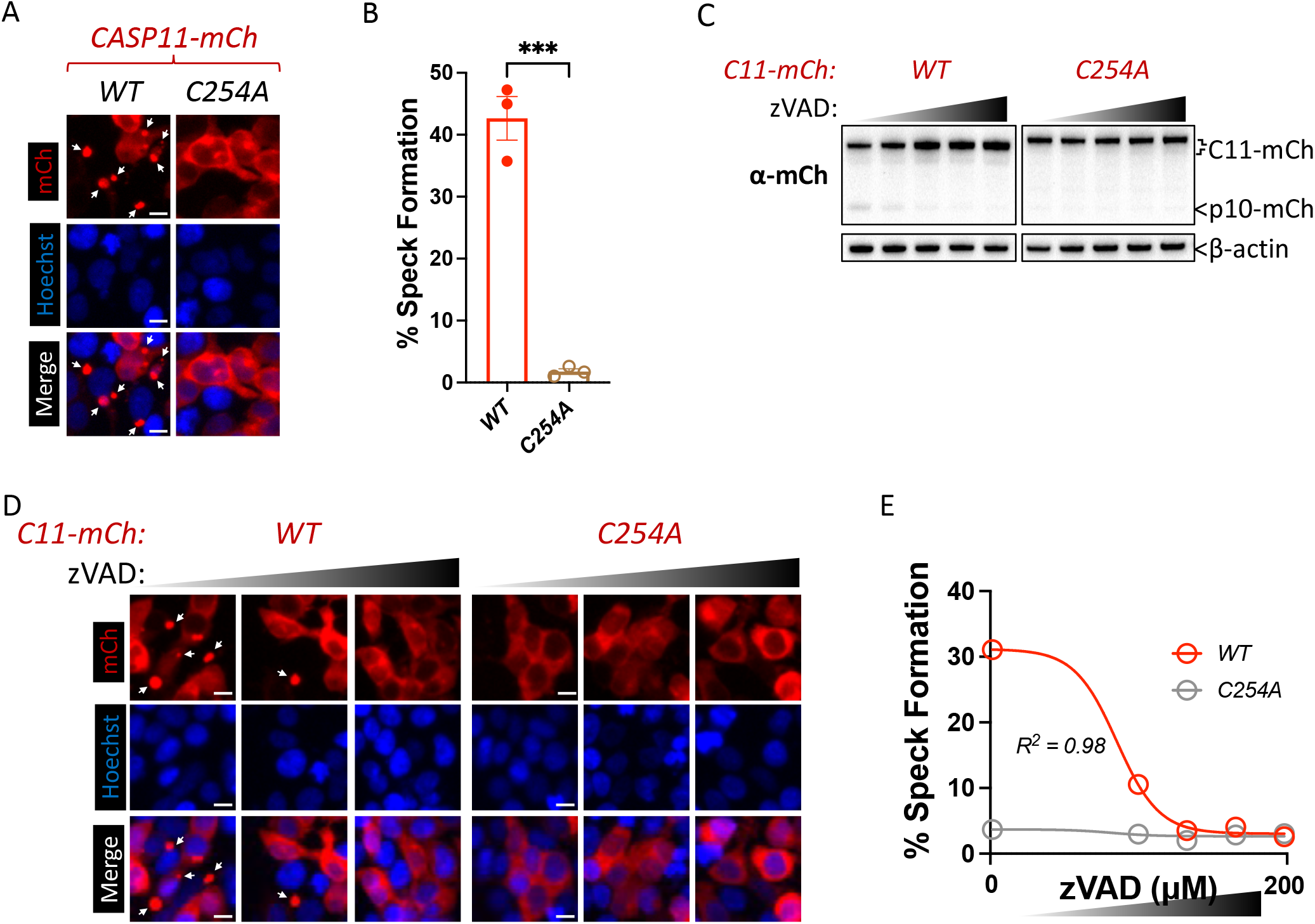
Caspase-11 catalytic activity is required for spontaneous caspase-11 oligomerization in HEK293T cells. (A) HEK293T cells were transfected with wild-type (*WT*) or catalytically inactive (*C254A*) Casp11-mCherry as described in Materials and Methods and imaged by fluorescence microscopy 18 h post transfection. Nuclei (blue) are stained with Hoechst, white arrows denote Casp11-mCherry specks. scale bar = 10 μm. (B) Speck formation in (A) was quantified as percentage of mCherry-expressing cells containing at least one speck. (C) HEK293T cells were transfected with Casp11-mCherry constructs as in (A) and 6 h post-transfection, cells were incubated with increasing amounts of pan-caspase inhibitor zVAD (0 – 200 μM; 2-fold increments). Whole cell lysates were isolated 12 hours post-transfection and immunoblotted for mCherry or β-actin as loading control as indicated. Cleaved p10-mCherry is denoted. (D) Casp11-mCherry speck formation was assayed in zVAD-treated cells by fluorescence microscopy as in (A). (E) Speck formation in (D) was quantified as percentage of mCherry-expressing cells containing at least one speck. Dose-response curves in were plotted using non-linear regression (Log_2_; three parameters). Error bars represent mean ± SEM of triplicate wells (800-900 cells per well); representative of 2±3 independent experiments. Data in (B) were analyzed by two-way ANOVA with Sidak’s multiple comparison test, ***p < 0.001.

### Caspase-11 processing at the interdomain linker is necessary but not sufficient for its oligomerization into specks

We next sought to understand the functional role of Casp11 catalytic activity in its assembly into oligomeric specks. Current established targets of Casp11 activity are itself, via cis- or trans-autoprocessing, and GSDMD. Since Casp11^WT^-mCherry spontaneously assembled into specks in 293T cells, which lack known substrates or signaling components for innate immune signaling pathways, we hypothesized that Casp11 autoprocessing might be important in its oligomerization. To test this, we generated Casp11^D285A^-mCherry, which contained a mutation in the interdomain linker (IDL) autoprocessing site. Like the catalytically inactive mutant, Casp11^D285A^-mCherry displayed impaired GSDMD cleavage and cell death as previously described (Lee *et al*., 2018; Ross *et al*., 2018), despite robust expression in HEK293T cells (**Extended Data Fig. 1C-E**). Intriguingly, Casp11 speck formation was significantly reduced in cells expressing Casp11^D285A^*-*mCherry in comparison to cells expressing Casp11^WT^-mCherry (**Fig. 3A, B**). To test whether autoprocessing at the IDL is *<u>sufficient</u>* to induce Casp11 oligomerization study this, we replaced the endogenous Casp11 cleavage sequence (_278_LPRN<u>MEAD</u>_285_) with the cleavage sequence for the tobacco etch virus (TEV) protease to generate a TEV-cleavable mCherry-tagged Casp11 construct: *Casp11-[_278_ENLYFQGA_285_]*-*mCherry* (Casp11^TEV^-mCherry) (**Fig. 3C**) (Chavarria-Smith et al., 2016; Oberst *et al*., 2010). We also mutated the catalytic cysteine on the TEV-cleavable construct, generating a Casp11 that could be inducibly cleaved with TEV but that was catalytically inactive (**Fig. 3C**). Co-expression of TEV protease with either WT- or C254A-Casp11^TEV^-mCherry in HEK293T cells resulted in dose-dependent Casp11 cleavage of both WT and C254A TEV constructs, as expected (**Fig. 3D**). Moreover, TEV protease induced Casp11 speck formation when co-expressed with Casp11^TEV^-mCherry, demonstrating that inducible TEV-dependent cleavage of Casp11 could rescue oligomerization of the autoprocessing-deficient mutant (**Fig. 3E, F**). Surprisingly, however, TEV protease-induced processing failed to rescue speck formation in the C254A Casp11^TEV^-mCherry mutant, suggesting that Casp11 catalytic activity also functions at an alternate site or that catalytic residue Cys-254 plays an additional role in inflammasome assembly (**Fig. 3E, F**). Altogether, these data indicate Casp11 autoprocessing at the IDL is necessary but not sufficient in the absence of catalytic activity to induce Casp11 oligomerization, suggesting that the catalytic cysteine contributes to noncanonical speck formation via additional mechanisms independent of IDL cleavage.

**Figure 3.**
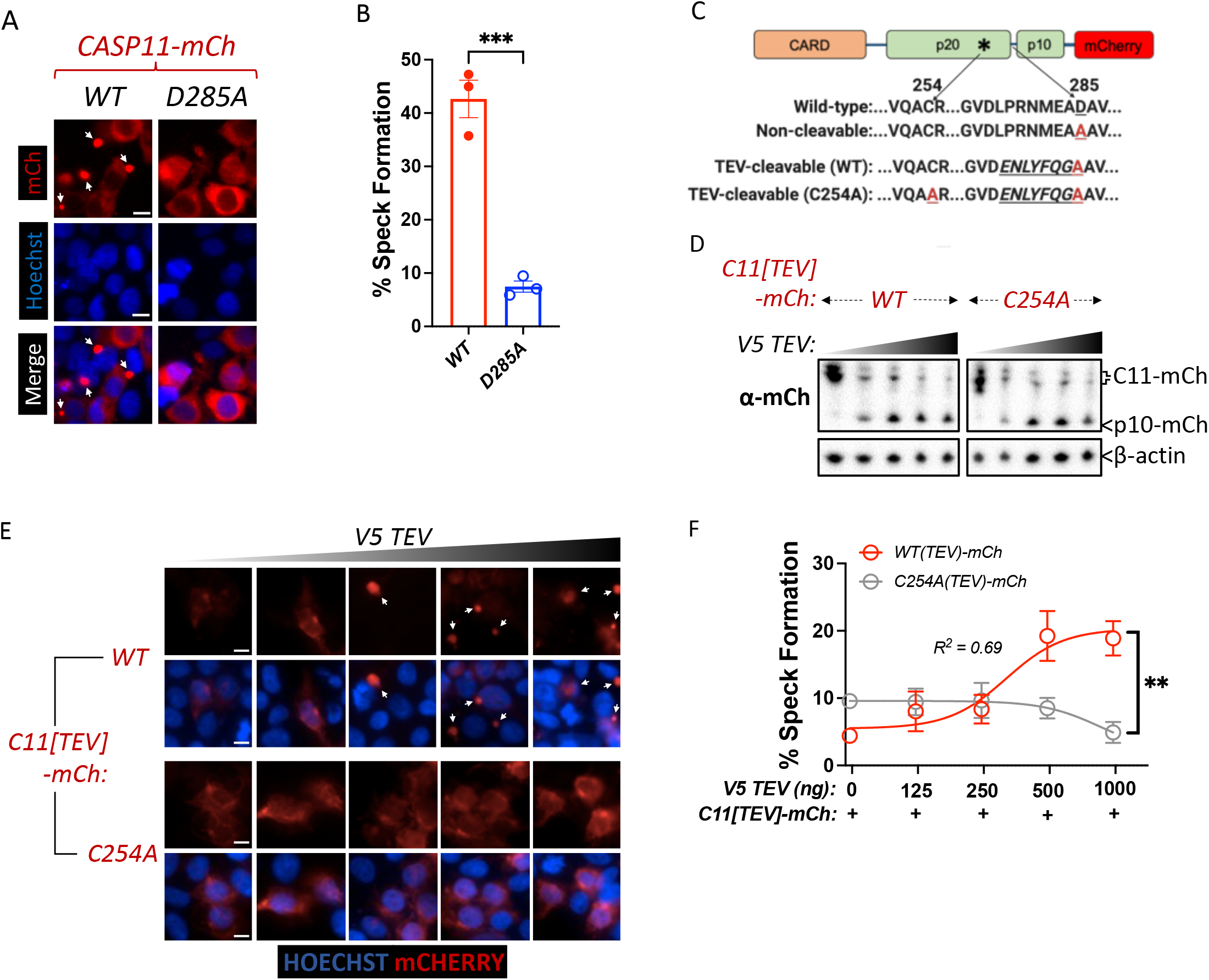
Caspase-11 processing at the interdomain linker is necessary but not sufficient for its oligomerization into specks. (A) HEK293T cells were transfected with wild-type (*WT*) or non-cleavable (*D285A*) Casp11-mCherry and imaged by fluorescence microscopy 18 h post-transfection. Nuclei (blue) are stained with Hoechst, white arrows denote Casp11 specks. scale bar = 10 μm. (B) Speck formation was quantified as percentage of mCherry-expressing cells containing at least one speck. (C) Schematic indicating Casp11-mCherry constructs that allow for inducible processing by tobacco etch viral (TEV) protease. The TEV protease consensus cleavage sequence (*<u>ENLYFQ/G</u>*) was engineered immediately upstream of cleavage-deficient Casp11-mCherry constructs with preserved (WT) or mutated (C254A) catalytic activity. (D) HEK293T cells were transfected with the indicated TEV-cleavable Casp11-mCherry constructs, together with increasing doses of V5-TEV protease (0 - 500 ng/well). Whole-cell lysates were harvested 12 hours post-transfection and immunoblotted for mCherry or β-actin (loading control). (E,F) 18 h post-transfection, cells were imaged by fluorescence microscopy and speck formation was quantified as in (A). Error bars represent mean ± SEM of triplicate wells (800-900 cells per well); representative of 2-3 independent experiments. Dose-response curves in (F) were plotted by non-linear regression (Log_2_; three parameters). Data were analyzed by two-way ANOVA with Sidak’s multiple comparison test, ***p <0.001.

### Wild-type caspase-11 rescues speck formation of catalytically inactive caspase-11, independently of inter-molecular ‘trans’ processing

Our data reveal that catalytic activity and autoprocessing within the IDL are critical for Casp11 speck formation. Casp11 autoprocessing can take place either in *trans* (intermolecular) or in *cis* (intramolecular). Given the ability of TEV-induced cleavage of an exogenous protease site in the IDL to induce speck formation, we anticipated that co-expression of wild-type unlabeled Casp11 might rescue oligomerization of a catalytic mutant Casp11^C254A^-mCherry. We therefore co-transfected fixed doses of labeled Casp11^WT^-mCherry or Casp11^C254A^-mCherry with increasing amounts of unlabeled wild-type Casp11 (**Fig. 4A**). Consistent with our prior studies, in the absence of co-transfected Casp11, Casp11^WT^-mCherry exhibited autoprocessing and speck formation that was absent in Casp11^C254A^-mCherry cells (**Fig. 4B**, left lanes, **4C**). Increasing levels of unlabeled Casp11 did not affect processing of Casp11^WT^-mCherry, though it did increase processing of Casp11^C254A^-mCherry (**Fig. 4B**), suggesting that while trans-processing of Casp11 could occur, processing of Casp11^WT^-mCherry was primarily due to cis processing. Notably, increasing levels of unlabeled Casp11 had a relatively limited effect on speck formation by Casp11^WT^-mCherry, except at the very highest dose (**Fig. 4C, D**). In contrast, expression of unlabeled Casp11 significantly increased speck formation by Casp11^C254A^-mCherry (**Fig. 4C, D**), demonstrating that indeed, a wild-type Casp11 could rescue oligomerization of catalytically inactive Casp11-mCherry species. Unexpectedly, however, and in contrast to the ability of TEV to mediate trans-processing of Casp11^WT^-mCherry, co-expression of untagged Casp11 did not lead to significant levels of Casp11^C254A^-mCherry processing, except at the highest concentrations (**Fig. 4B**). These data indicated that Casp11 does not induce significant levels of trans-processing in the course of oligomerization into specks. Indeed, additional mutation of the IDL autoprocessing site of the Casp11^C254A^-mCherry did not affect its ability to form specks in the presence of co-transfected WT unlabeled Casp11 (**Fig. 4G, H**). Altogether, these studies demonstrate that both catalytically inactive, as well as inactive-uncleavable, Casp11 can be recruited into oligomeric inflammasome complexes in the presence of catalytically active Casp11, which must undergo intra-molecular processing in order to induce formation of a Casp11 SMOC.

**Figure 4.**
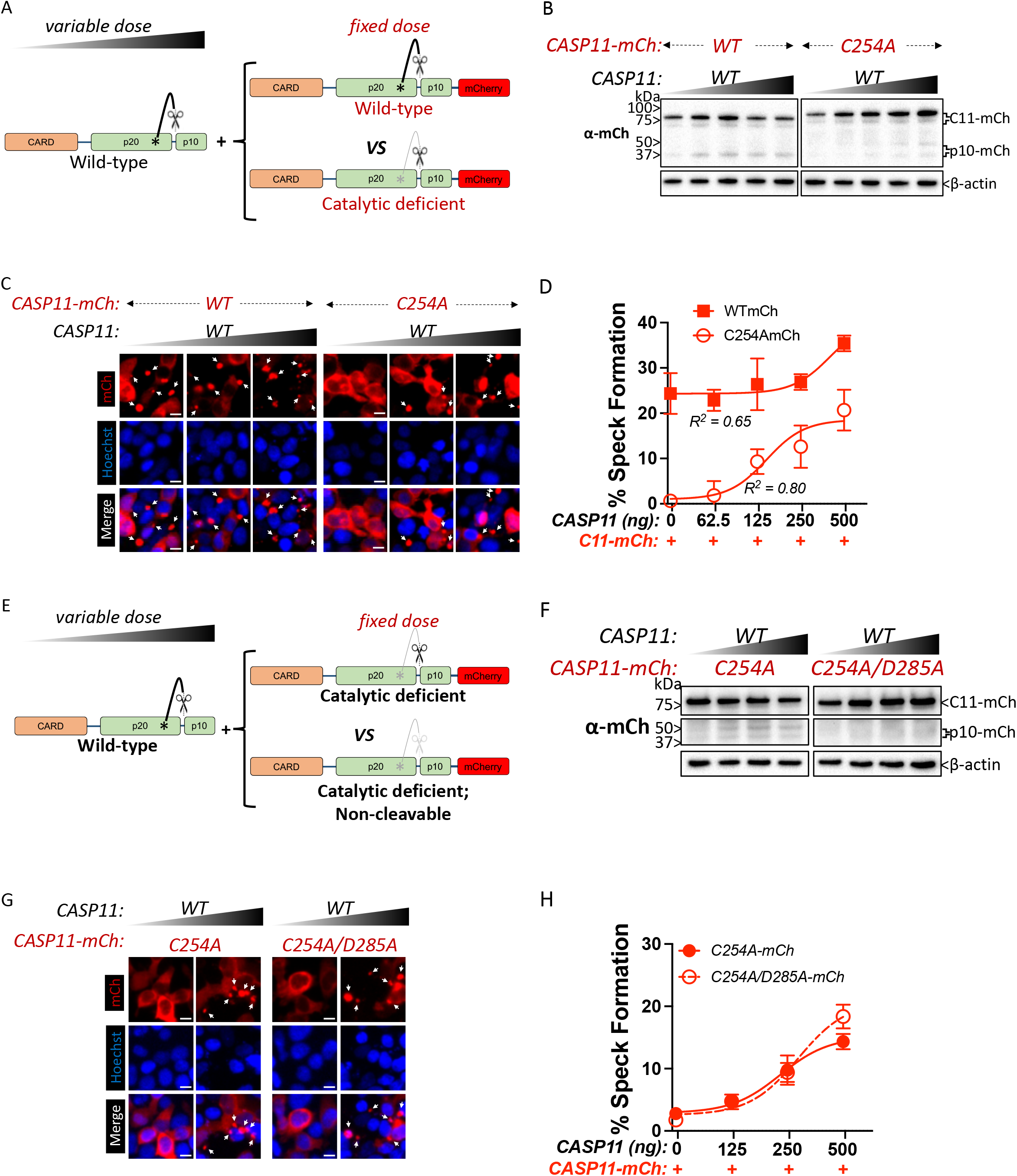
Wild-type caspase-11 rescues speck formation of catalytically inactive caspase-11, independently of inter-molecular ‘trans’ processing. (A, E) Schematic diagram indicating co-transfection combinations used in (B-F). Untagged full-length wild-type (*WT*) caspase-11 gene constructs were transfected with increasing doses together with a fixed amount of indicated mCherry-tagged WT, C254A, or C254A/D254A double-mutant Casp11, as indicated. (B) 12 h post-transfection, whole-cell lysates were harvested and immunoblotted for mCherry or β-actin as a loading control. (C) 18 h following transfection, the cells were imaged by fluorescence microscopy. Nuclei (blue) are stained with Hoechst, white arrows represent Casp11 oligomers (specks), scale bar = 10 μm. (D) Speck formation was quantified as percentage of mCherry-expressing cells containing at least one speck. (F) Whole cell lysates of indicated transfected cells was assayed as in (B). (G) Cells transfected with indicated constructs were imaged as in (C) and speck formation with respect to increasing levels of WT Casp11 was plotted as in (D). Error bars indicate mean ± SEM of triplicate wells (800-900 cells per well); representative of 2-3 independent experiments. Dose-response curves in (D, H) were plotted by non-linear regression (Log_2_; three parameters).

### Caspase-11 cis autoprocessing mediates noncanonical inflammasome assembly

The above observations imply that while catalytically deficient Casp11 species lack the ability to oligomerize autonomously, co-expression with a catalytically competent partner can rescue this oligomerization independently of trans-processing. To directly test the possibility that this rescue is driven by cis autoprocessing of the catalytically competent Casp11 species, we co-transfected labeled Casp11^C254A^-mCherry with either catalytically inactive Casp11^C254A^ or non-cleavable Casp11^D285A^ unlabeled Casp11 species (**Fig. 5A**). A prediction of this model is that catalytic activity and autoprocessing of unlabeled Casp11 should be required to rescue speck formation by Casp11^C254A^-mCherry. As expected, WT Casp11 could undergo autoprocessing, which was absent in cells transfected with either the catalytic or autoprocessing mutants (**Fig. 5B**). Critically, WT unlabeled Casp11 again rescued speck formation by catalytically inactive Casp11-mCherry (**Fig. 5C, D**). Importantly however, neither unlabeled Casp11^C254A^ nor Casp11^D285A^ could restore speck formation for Casp11^C254A^-mCherry (**5C, D**). These findings suggest that cis autoprocessing of Casp11 results in assembly of a functional oligomeric complex to which the labeled Casp11 reporter mutants can be recruited (**Fig. 5E**). Altogether, these findings demonstrate a key role for Casp11 catalytic activity and cis autoprocessing in oligomerization and higher order assembly of the Casp11 inflammasome. Furthermore, our findings indicate that these activities occur upstream of higher order Casp11 inflammasome assembly, rather than occurring as a terminal consequence of inflammasome assembly.

**Figure 5.**
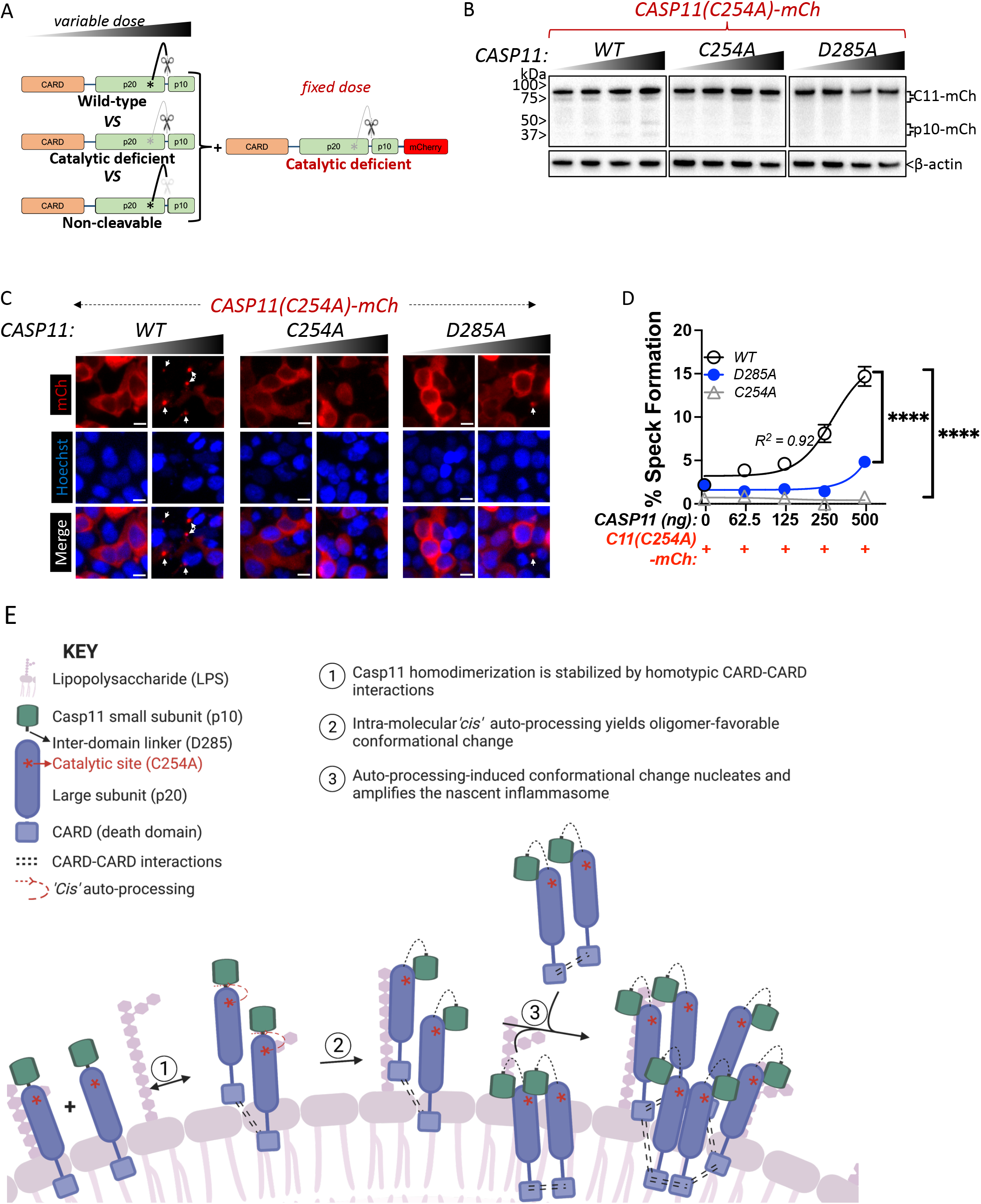
Caspase-11 cis autoprocessing mediates noncanonical inflammasome assembly. (A) Schematic diagram indicating co-transfection combinations used in (B-D). Unlabeled wild-type (*WT)*, catalytic-deficient (*C254A*) or non-cleavable (*D285A*) caspase-11 constructs were transfected into HEK293T cells at increasing doses together with a fixed dose of catalytic-deficient (C254A) mCherry-tagged caspase-11. (B) 12 h post-transfection, whole-cell lysates were harvested and immunoblotted for mCherry or β-actin as loading control. (C) 18 h following transfection, cells were imaged by fluorescence microscopy. Nuclei (blue) are stained with Hoechst, white arrows denote caspase-11 specks, scale bar = 10 μm. (D) Speck formation was quantified as percentage of mCherry-expressing cells containing at least one speck. Error bars = mean ± SEM of triplicate wells (800-900 cells per well); representative of 2-3 independent experiments. (E) Proposed mechanism of caspase11 inflammasome assembly (created with BioRender.com). Dose-response curves were plotted by non-linear regression (Log_2_; three parameters). Two-way ANOVA (highest doses) with Sidak’s multiple comparison test, ****p <0.0001.

## DISCUSSION

Casp11 is a direct sensor of cytosolic LPS and plays a vital role in host defense against infection (Aachoui et al., 2013). Dysregulation of Casp11 or its human orthologs, Casp4 and Casp5, leads to severe inflammatory disorders such as endotoxic shock and experimental autoimmune encephalomyelitis (Hagar *et al*., 2013; Kajiwara et al., 2014; Kayagaki *et al*., 2013; Napier et al., 2016). LPS binds to the CARD of Casp11 and promotes its assembly into a large oligomeric complex termed the noncanonical inflammasome (Shi *et al*., 2014). Given their critical roles in innate immune responses to disease states resulting from infection, auto-inflammatory disease, and sepsis, there is substantial interest in understanding how assembly of such higher order oligomeric complexes, also known as supramolecular organizing centers, or SMOCs, is regulated (Kagan *et al*., 2014).

Dimerization of Casp11 is sufficient to induce its protease activity to undergo autoprocessing, which is required for its ability to cleave GSDMD, suggesting that activation of the fully competent oligomeric complex occurs subsequent to initial clustering of Casp11 (Ross *et al*., 2018). Genetic studies further reveal that both catalytic activity at C254 and autoprocessing at D285 within the interdomain linker region between the two enzymatic subunits (IDL), are required for Casp11-mediated processing of GSDMD and inflammatory function (Lee *et al*., 2018). However, the requirement for Casp11 autoprocessing for its ability to cleave GSDMD even when dimerized and catalytically active (Ross *et al*., 2018) implies that dimerization alone is not sufficient for complete activation of Casp11. This requirement for both dimerization and autoprocessing in Casp11 activation is reminiscent of the requirement for dimerization and autoprocessing for full activity of Casp8 and human Casp1 (Ball *et al*., 2020; Oberst *et al*., 2010). These findings suggest that initial dimerization and subsequent oligomerization might be two distinctly regulated steps in activating the fully competent Casp11 noncanonical inflammasome. Since Casp11 catalytic activity lies at the center of both its autoprocessing and GSDMD processing, we sought to understand how this enzymatic activity might contribute to assembly of Casp11 complexes.

Using a Casp11 reporter fusion protein to track the subcellular localization of Casp11 in individual cells via fluorescence microscopy, we found that cytosolic LPS stimulation induced assembly of perinuclear Casp11 clusters in BMDMs, similar to ASC inflammasome specks observed in cells expressing the ASC-Citrine fusion protein (Stutz *et al*., 2013; Tzeng *et al*., 2016). Surprisingly, mutating the catalytic site on Casp11-mCherry revealed that Casp11 catalytic activity is required to form Casp11 specks in macrophages in response to cytosolic LPS. This implies that CARD-LPS binding is insufficient to mediate Casp11 SMOC formation. Likewise, ectopic expression of Casp11-mCherry fusion proteins in HEK293T cells, along with corresponding catalytically inactive and autoprocessing deficient mutants, showed that both Casp11 catalytic activity and autoprocessing are required for spontaneous recruitment of Casp11 into oligomeric complexes (**Figs. 2A-B, 3A-B**). Collectively, our findings suggest that Casp11 enzymatic activity and autoprocessing function through a feed-forward mechanism to promote assembly of the fully active noncanonical inflammasome.

HEK293T cells have been used extensively to dissect the molecular and cellular determinants of inflammasome assembly innate immune signaling (Evavold et al., 2021; Kayagaki *et al*., 2013; Sandstrom et al., 2019; Shi *et al*., 2014; Tenthorey *et al*., 2014). Importantly, the requirement for catalytic activity in speck formation was seen both in 293T cells and in LPS-inducible speck formation in primary BMDMs. Collectively, these studies demonstrate that spontaneous speck assembly in HEK293T cells follows similar rules to LPS-induced speck assembly in macrophages, and that the HEK293T system can be used to understand how the noncanonical inflammasome is assembled.

Casp11 processing by the exogenous TEV protease restored speck formation of Casp11 constructs whose endogenous IDL cleavage site was replaced with a TEV protease site, indicating that Casp11 processing is sufficient for speck formation (**Fig. 3C-F**). Surprisingly, the Casp11 catalytic Cys-254 was still required for speck assembly of Casp11, as C254A Casp11^TEV^-mCherry constructs failed to form specks despite undergoing TEV processing (**Fig. 3C-F**). This finding suggests that Casp11 must either undergo processing at an additional site, such as Asp-80 located at the CARD domain linker (CDL) or that catalytic residue Cys-254 plays an alternate unknown role for speck assembly. Notably, the CDL of Casp1 is a negative regulator of Casp1 activity, and the CDL of Casp11 was previously suggested to have no impact on Casp11 activity (REFS) (Boucher *et al*., 2018; Ross *et al*., 2018). Thus the alternate target of Casp11 outside of the IDL processing site is currently not clear. Importantly, C254A mutant Casp11 was capable of binding to both WT Casp11, and C254A Casp11, indicating that loss of binding per se was not the basis for the inability of Casp11 C254A to undergo oligomerization (**Extended Data Fig. 3**).

Regardless, our studies demonstrate that in the presence of a functional catalytic site, autoprocessing is sufficient to form the Casp11 speck, implicating autoprocessing in noncanonical inflammasome assembly.

Our studies find that wild-type unlabeled Casp11 rescued speck formation of catalytically inactive Casp11-mCherry (**Figs. 4A-D**). Given that TEV protease was unable to perform this function despite robust processing of Casp11-mCherry, this suggested that rather than providing a cleavage function, unlabeled Casp11 was recruiting inactive Casp11 into a complex formed by catalytically competent Casp11. Indeed, labeled Casp11 did not require its catalytic activity or processing to form specks in the presence of unlabeled wild-type Casp11 (**Fig. 4E-H**), whereas both catalytic activity and processing were necessary for unlabeled Casp11 to induce speck formation of catalytically inactive Casp11-mCherry (**Fig. 5A-D**). Collectively, these data imply that intra-molecular cis-autoprocessing, rather than trans processing drives assembly of higher order Casp11 oligomeric complexes, supporting a cooperative feed-forward mechanism by which Casp11 dimerization and cis processing, likely induced by LPS binding, propagates the assembly of a fully active Casp11 SMOC. Notably, recent cryo-EM analyses revealed that human NLRP1 and CARD8 propagate their inflammasomes through a similar autoprocessing-dependent model (Gong et al., 2021). Likewise, cooperative assembly has been proposed for Casp8 assembly onto the Death Inducing Signaling Complex (DISC) (Fox et al., 2021), suggesting that this mechanism may generally apply to other oligomeric caspase complexes.

Catalytically inactive recombinant Casp11 purified from insect cells undergoes aggregation into higher molecular weight complexes in response to treatment with cell lysates from gram-negative bacteria or purified LPS (Shi *et al*., 2014). These studies utilized highly purified components to define mechanics of Casp11 assembly *in vitro*. Here, we utilized a combination of genetic and pharmacologic approaches to visualize the behavior of Casp11 within individual cells. Based on the behavior of analogous ASC specks, we interpret the Casp11 specks that we observe to be higher order oligomeric Casp11 structures. It is possible that regulatory mechanisms within the cytosol of intact cells limit the aggregation of catalytically inactive or non-cleavable Casp11. A previous study found that EGFP-tagged Casp11 exhibited speck formation in WT and catalytically inactive Casp11-expressing 293T cells (Liu et al., 2020). Notably, these Casp11-EGFP specks appear smaller (<1–2 μm) and less defined than the spherical perinuclear specks (2-5 μm) we observed in mCherry or Citrine fusion-expressing cells. Moreover, our observations in HEK293T cells recapitulate the endogenous behavior of the Casp11 SMOC in macrophages, as catalytic activity is required for LPS-induced Casp11 assembly in macrophages as well as spontaneous Casp11 speck assembly in 293T cells.

While Guanylate-Binding Proteins (GBPs) act as important facilitators of Casp11 activation (Man et al., 2016; Meunier et al., 2014; Pilla et al., 2014; Wandel et al., 2020), Casp11 contains all the key features necessary to form a noncanonical inflammasome, as it contains the properties of an inflammsome sensor, adaptor, and effector within a single protein. Interestingly, while the Casp11 CARD is sufficient to binds LPS (Shi *et al*., 2014), we find that this domain is not sufficient for spontaneous Casp11 oligomerization in HEK293 cells, further supporting the finding that critical oligomerization functions are provided by catalytic activity and autoprocessing. Precisely how autoprocessing mediates Casp11 higher order oligomerization remains to be determined. Initial Casp11 homodimerization and autoprocessing may induce a conformation change that enhances the binding affinity between different homodimers, resulting in a shift in the equilibrium to favor sequential recruitment of further monomers to propagate the fully assembled the oligomeric complex. Future studies on the oligomeric structure of Casp11 are likely to reveal important information about how the individual Casp11 monomers interact with one another, and how CARD-LPS interactions facilitate Casp11 oligomerization. Our findings provide new insight into the assembly of higher order caspase complexes, and implicate catalytic activity and autoprocessing in the assembly of the noncanonical inflammasome.

## MATERIALS AND METHODS

### Cell culture

HEK293T cells were purchased from ATCC, grown in complete DMEM (supplemented with 10% v/v FBS, 10 mM HEPES, 10 mM Sodium pyruvate, 1% Penicillin/Streptomycin), and maintained in a 37°C incubator with 5% CO_2_. Murine bone marrow progenitors were harvested from femurs and hip bones of 8-12 week-old *C57BL/6* or *Casp11*^-/-^ mice and differentiated into bone marrow-derived macrophages (BMDMs) using 30% L929-conditioned complete DMEM for 7 days as previously described (Bjanes et al., 2021).

### Cloning

The Casp11 coding sequence was fused to mCherry on its C-terminus, yielding the fusion reporter construct *CASP11-mCHERRY* on pTwist Lenti-SFFV-WPRE lentiviral vector (Twist Biosciences) or mammalian expression vector pReceiever-M56 (Genecopoeia). Caspase-11 catalytic (C254A) and cleavage (D285A) mutants were generated by site-directed mutagenesis (Q5 SDM Kit, New England BioLabs; #E0554S) of the wild-type parent vector according to manufacturer’s instructions. To generate TEV-cleavable constructs, the TEV protease consensus cleavage sequence (*<u>ENLYFQ/G</u>*) was engineered immediately upstream of cleavage-deficient (D285A) Casp11-mCherry construct on the on pTwist Lenti-SFFV-WPRE lentiviral vector backbone. Catalytical site mutants (C254A) were generated as above. All constructs were confirmed by sequencing prior to experimentation.

### HEK293T Transient transfections

Mammalian expression plasmids containing indicated DNA constructs were transfected into HEK293T cells using the transfection agent polyethylenimine (PEI; Sigma-Aldrich) at 1:1 ratio (w/w DNA:PEI) in Opti-MEM (Gibco). Media was changed to complete DMEM (10% v/v FBS) after 4-6 hours, and cells were incubated for subsequent times as indicated in figure legends in a humidified incubator at 37°C and 5% CO_2_ prior to subsequent analysis.

### Lentiviral transduction

Lentiviral plasmids containing *WT* or *C254A CASP11-mCHERRY*, along with pVSV-G and psPAX2 packaging plasmids, were transfected into HEK293T cells using PEI (Sigma-Aldrich) at 1:1 ratio (w/w DNA:PEI) in low-serum media (complete DMEM with 2% v/v FBS). After 12 hours, media was changed to complete DMEM (10% v/v FBS) and cells were incubated for another 48 h at 37°C and 5% CO_2_ for virus production. The resulting lentiviral supernatants were filtered (0.45 μm), and concentrated with Lenti-X concentrator (Takara Bio) according to manufacturer’s instructions. Viral prep was resuspended in 30% L929-supplemented complete DMEM (mac media), supplemented with 10 μg/ml polybrene, and used to spin-infect Day 2 *Casp11*^-/-^ bone marrow progenitors (1750 RPM for 90 min at 22°C). Progenitors were replaced in 37°C incubator and allowed to differentiate until day 7 as described above.

### Cytosolic LPS Delivery, LDH Cytotoxicity Assay & ELISA

Following differentiation into mature macrophages, primary murine BMDMs were seeded in TC-treated 96-well plates (6.0 × 10^4^ cells/well) and left to adhere in 37°C incubator overnight. On day of LPS transfection, cells were primed with bacterial membrane mimetic Pam3CSK4 (400 ng/mL; Invivogen) for 4 h. LPS from *S. enterica* serotype Minnesota (Sigma) was then packaged into stable complexes using FuGENE HD (Promega; E2311) and added to cells in Opti-MEM (Gibco) as previously described (Hagar *et al*., 2013; Harberts et al., 2022). 16 h following LPS transfection, supernatants were harvested and assayed for lactate dehydrogenase (LDH) release as a read-out for pyroptosis, as previously described (Mariathasan et al., 2004; Rayamajhi et al., 2013). Briefly, at the appropriate timepoints, plates were spun down (250 ×g) for 5 min to rid supernatants of cell debris. Supernatants (50 μL) were then combined with an equivalent volume of LDH reaction buffer/substrate mix (Takara Bio Inc.) in a clear-bottom 96-well plate. After ~20 mins at room temp, absorbance was read on a spectrophotometer (495 nm) and normalized to mock-transfected cells (min cell lysis; negative control) and cells treated with 1% TritonX-100 (max cell lysis; positive control). To assess IL-1β release, supernatants were diluted 4-fold and applied to Immulon ELISA plates (ImmunoChemistry Technologies) pre-coated with anti-IL-1β capture antibody (eBioscience). Following blocking (1% BSA in 1× PBS), plates were incubated with biotin-linked secondary antibody, followed by horseradish peroxidase-conjugated streptavidin. As read-out for IL-1β levels, peroxidase enzymatic activity was determined by exposure to o-phenylenediamine hydrochloride (Sigma) in citric acid buffer. Reactions were stopped with sulfuric acid and absorbance values were read at 490 nm, normalized to mock-transfected cells (negative control).

### Cell Viability Assay

Viability of HEK293T cells co-transfected with GSDMD and various caspase-11 expression constructs was determined using the CellTiter-Glo 2.0 Assay Kit (CTG2.0; Promega) according to manufacturer’s instructions. Briefly, HEK293T cells were seeded in poly-L-lysine-coated (0.1 mg/mL; Sigma-Aldrich) F-bottom 96-well plates and allowed to adhere overnight in complete DMEM. The next day, mammalian expression plasmids containing indicated mCherry-tagged or -untagged caspase-11 DNA constructs were co-transfected into the cells at increasing doses (0 – 250 ng per well) together with a fixed dose of GSDMD (50 ng), using the transfection agent polyethylenimine (PEI; Sigma-Aldrich) at 1:1 ratio (w/w DNA:PEI). After 14 h, cells were lysed with CTG2.0 reagent mix and incubated in the dark at 37 °C for 30 mins. Luminescence was read on a luminometer, and values were normalized to cells treated with 1% TritonX-100 (min cell viability; negative control) and mock-transfected cells (max cell viability).

### Immunoblotting

BMDMs were seeded in clear TC-treated 96-well plates (6.0 × 10^4^ cells/well) and transfected with LPS in the absence or increasing doses of zVAD-fmk (caspase-11 inhibitor), as indicated. HEK293T cells were seeded in poly-L-lysine-coated TC-treated 24-well plates (2.0 × 10^5^ cells/well) and transiently transfected with appropriate gene constructs as described above. At the indicated timepoints, whole-cell extracts (XT) and/or supernatants (sup) were harvested and immunoblotted for various proteins as previously described (Bjanes *et al*., 2021; Harberts *et al*., 2022). In brief, plates were centrifuged (250 ×g) to rid supernatants of cell debris. Proteins in supernatants were then precipitated by incubating with 0.61 N trichloroacetic acid (Sigma-Aldrich) plus 1× protease inhibitory cocktail (PIC; Sigma-Aldrich) for at least 1 h on ice. Precipitates were washed with acetone by centrifugation (×3) at 4°C, and resuspended in protein sample buffer (125 mM Tris, 10% SDS, 50% glycerol, 0.06% bromophenol blue, 1% β-mercaptoethanol, 50 mM dithiothreitol). Lysates were harvested in lysis buffer (20 mM HEPES, 150 mM NaCl, 10% glycerol, 1% Triton X-100, 1 mM EDTA, pH 7.5) supplemented with 1× PIC and 1× protein sample buffer and rocked gently for 10 min at 4°C. Protein samples from lysates and supernatants were then prepared for SDS-PAGE by boiling, centrifugation (table top max speed; 5 mins) before they were run on 4-12% polyacrylamide gels (Invitrogen). Proteins were transferred to polyvinylidene difluoride (PVDF) membranes and immunoblotted with the following antibodies: caspase-11 (1:1,000; Novus Biologicals 17D9), mCherry (1:1000; Abcam EPR20579), or GSDMD (1:500; Abcam EPR19828). β-actin (1:2,500; Sigma-Aldrich AC74) was used as a loading control. Blots were then incubated in species-specific, horseradish peroxidase-conjugated secondary antibodies (1:2,500) and imaged by chemiluminescence using Pierce SuperSignal West Femto maximum sensitivity substrate (Thermo Scientific) according to the manufacturer’s instructions.

### Casp11 immunoprecipitation

HEK 293T cells were transiently transfected with the indicated FLAG-tagged construct for 48 hours. Collected cell pellet samples were lysed by sonication and clarified by centrifugation at 20,000 × *g* for 5 minutes at 4 °C. Aliquots from the clarified lysate samples were incubated with ANTI-FLAG M2 affinity beads (Millipore Sigma) in Pierce Micro-Spin columns (ThermoFisher Scientific) at 4 °C overnight. The Micro-Spin samples were then washed three times with one column volume of PBS, and proteins eluted with 3x-FLAG peptide followed by Western blot analysis. Aliquots of the remaining lysate was used to prepare input samples for Western analysis.

### Fluorescence and confocal microscopy

BMDMs or HEK293T cells were seeded on round, poly-L-lysine-coated #1.5H glass coverslips (Thorlabs, #CG15NH) and allowed to adhere overnight. Cells were then transfected with LPS (BMDMs) or indicated DNA constructs (HEK293Ts). At the indicated timepoints, cells were washed 1× with PBS and fixed with 4% paraformaldehyde. Following nuclear counterstain with Hoechst 33342 (1 μg/mL; Thermo Fisher #62249), cells were mounted on glass slides with Fluoromount-G (SouthernBiotech; 0100-01) and dried overnight. Slides were imaged on brightfield, Cy3 (red) and DAPI (blue) fluorescence channels using a Dmi8 inverted wide-field apparatus at 20X objective (Leica Biosystems). For confocal microscopy, slides were imaged at a single z-plane per field with lasers optimized for Cy5 (far-red), Citrine (yellow) and cell-tracker violet (blue) emission spectra, through 63X objective.

### Image Quantification and Analysis

Each experiment was conducted in three technical replicates. Within each replicate, 4-6 frames were analyzed for an average of 100-150 cells (BMDMs) or 800-900 cells (HEK293T cells) per well. A speck was defined as a distinct high-fluorescent perinuclear cluster of mCherry or citrine. Speck formation frequency was determined as the percentage of mCherry-expressing (red) or citrine-expressing (yellow) cells that contained one or more specks, using custom macros from ImageJ (NIH) and LAS X (Leica Biosystems).

### Statistical Analysis

Data were graphed and analyzed using GraphPad Prism 9 (San Diego, CA). Mean values (± SEM) were compared across triplicate conditions and P values were determined using one sample t-test, one-way or two-way analysis of variance (ANOVA) with Sidak’s multiple comparison. Dose-response curves were plotted using non-linear regression (Log_2_ scale; three parameters).

**Extended Data, Figure 1. Casp11-mCherry maintains enzymatic function in HEK293T cells.** (A) Schematic representation of Casp11 fluorescent reporter constructs used in this study indicating mCherry fused to the C-terminus of wild-type (WT), catalytically inactive (C254A) or non-cleavable (D285A) *Casp11*. (B) Indicated *Casp11-mCherry* expression plasmids were transfected into HEK293T cells. Cell lysates were immunoblotted for mCherry, Casp11, and β-actin (loading control) ten hours post-transfection. (C-E) Gasdermin-D (GSDMD) expression plasmid was co-transfected with increasing doses of indicated Casp11 gene construct in HEK293T cells. WT untagged *CASP11* was included as positive control. After 12 h (C) or 16 h (D, E), Casp11 activity was determined by (C) immunoblotting for GSDMD processing in supernatants (sup) and whole-cell lysates (XT), (D) cytotoxicity, and (E) cell viability as indicated in materials and methods. Error bars indicate mean ± SEM of triplicate wells; representative of 2-3 independent experiments. Dose-response curves were plotted using non-linear regression (Log2; three parameters). Two-way ANOVA with Sidak’s multiple comparison test (highest doses), *p < 0.05, ****p <0.0001.

**Extended Data, Figure 2. Casp11 catalytic activity mediates GSDMD cleavage, pyroptosis and IL-1β release in response to intracellular LPS in primary BMDMs.** Wild-type (*B6*) or *Casp11*-/-primary BMDMs were primed with Pam3CSK4 for 4 h, followed by transfection with the indicated concentrations of LPS from *S. enterica serotype Minnesota*. To inhibit Casp11 activity, cells were incubated with indicated concentrations of zVAD beginning 30 mins before (and lasting through) LPS transfection. After 16 h, pyroptosis was measured by (A) assessing supernatants for lactate dehydrogenase (LDH) release (normalized to maximum cell lysis by TritonX-100), (B) analyzing IL-1β release in supernatants by ELISA, and (C) immunoblotting for gasdermin D (GSDMD) cleavage in supernatants (sup) and whole-cell lysates (XT). β-actin is indicated as loading control. Error bars = mean ± SEM of triplicate wells; representative of 2-3 independent experiments. *p < 0.05, **p < 0.01, ***p <0.001, ****p < 0.0001, ns not significant. Two-way ANOVA with Sidak’s multiple comparison test.

**Extended Data, Figure 3. Catalytic activity is not required for caspase-11 intermolecular interactions.** (A) Tagged caspase-11 expression used for FLAG-based co-immunoprecipitation. (B)HEK293T cells were transiently transfected with 2X-FLAG-tagged wild-type (*WT*) or catalytically inactive *Casp11* expression plasmids alongside *WT* or *C254A* mCherry-tagged *Casp11* (5 μg). 48 hours post-transfection, whole-cell lysates were immunoprecipitated by anti-FLAG antibodies as described in Materials and Methods, and immunoblotted for mCherry, FLAG or GAPDH as a loading control, as indicated.

**Extended Data, Figure 4. Casp11 CARD is not sufficient to mediate Casp11 oligomerization in HEK293T cells.** (A) Schematic diagram describing fluorescent reporter constructs used in (B, C). (B) The fluorescent reporter citrine was fused to the C-terminus of wild-type (*WT*) or CARD-only *Casp11* and HEK293T cells were transfected with indicated constructs and imaged by confocal microscopy 18 hours post-transfection. Nuclei (magenta) are stained with DRAQ7, cytosolic content (blue) was stained with cell tracer violet. (C) Citrine-based speck formation (yellow) was quantified as percentage of overall citrine-expressing cells that contained at least one speck. Error bars represent mean ± SEM of triplicate wells (800-900 cells per well); a single experiment was performed. Two-way ANOVA with Sidak’s multiple comparison test, *p <0.05, **p < 0.01, ***p < 0.001, ns = not significant.

## Acknowledgments

We are grateful to members of the Brodsky, Shin and Taabazuing labs for scientific discussion. We would like to thank the Herbert and Vaughan labs for use of lab equipment and the Leica Dmi8 inverted wide-field microscope. This work was supported by grants R01AI128530, R01139102A1, and an awards from the Mark Foundation (IEB), R01AI118861 and R01AI123243 (SS) and the Burroughs Welcome Fund Investigator in the Pathogenesis of Infectious Disease Award (IEB and SS); and UNCF/BMS EE Just Early Career Investigator Award and NIH R00 Career Transition Award Grant# 4R00AI148598-03 (CYT).

