## Supplementary figures and images for "Noncanonical inflammasome assembly requires caspase-11 catalytic activity and intra-molecular autoprocessing"

### Extended Data Figures

#### Extended Data Figure 1

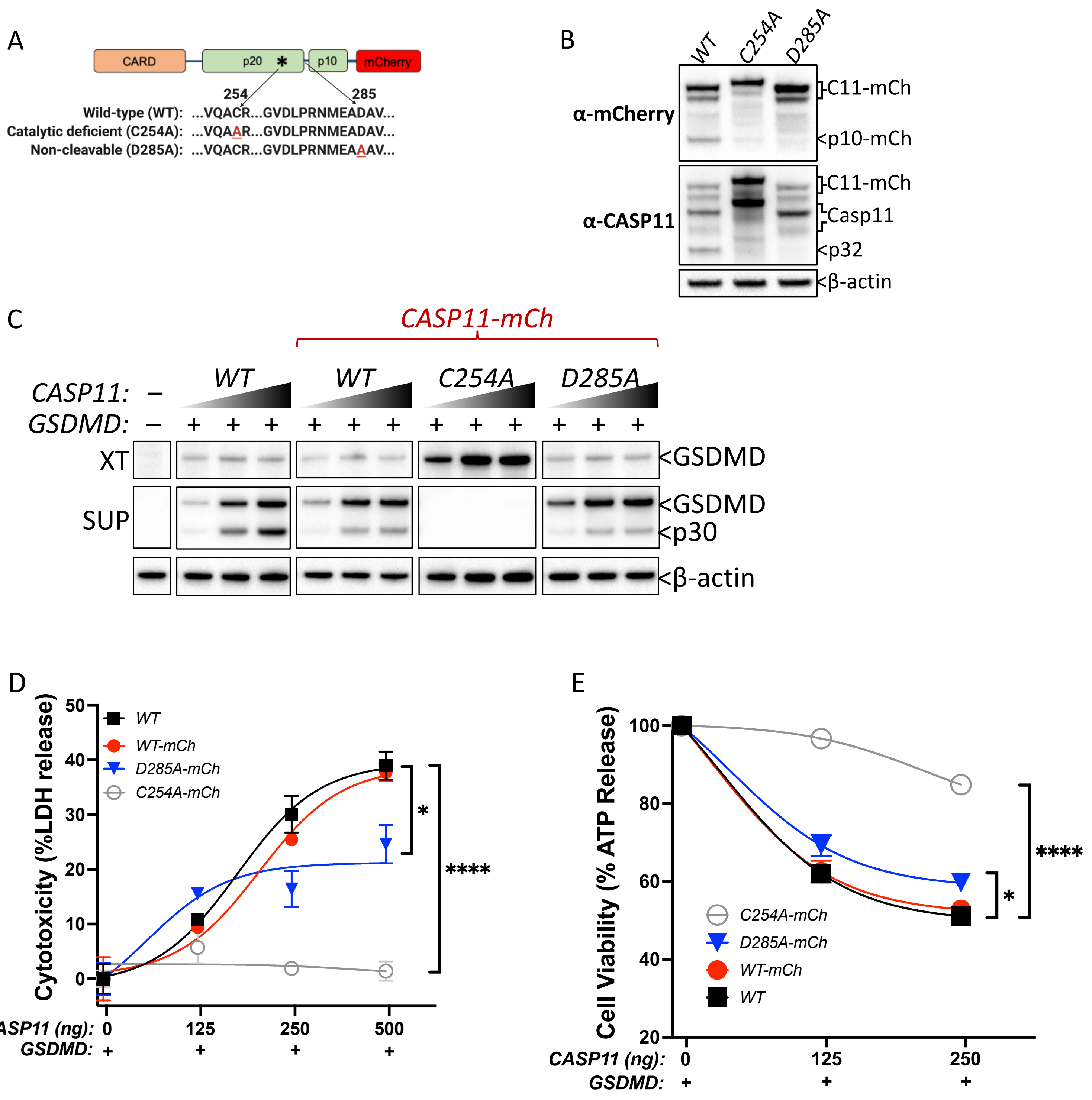

#### Extended Data Figure 2

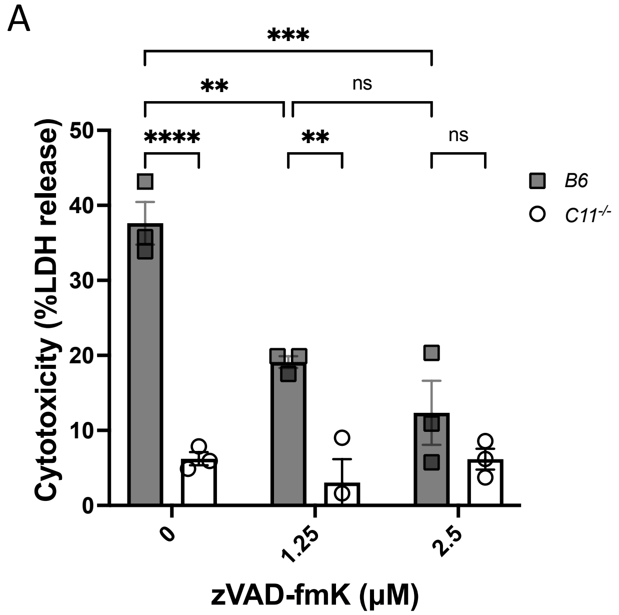


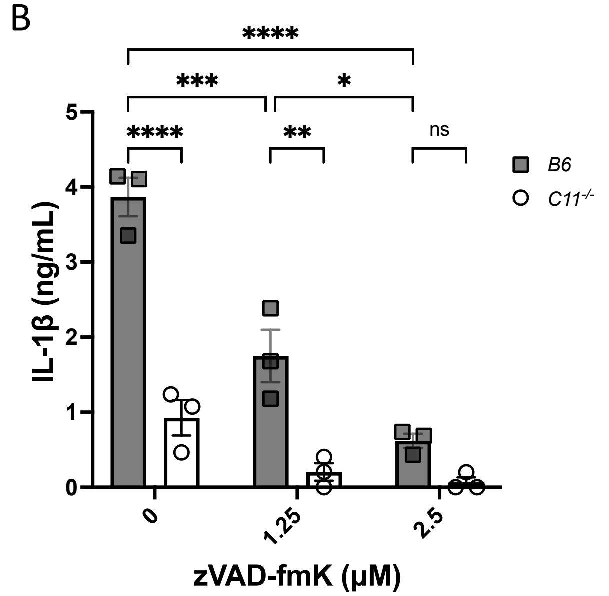


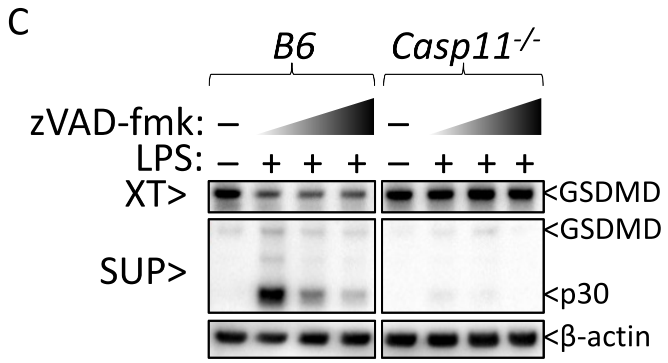

#### Extended Data Figure 3

A

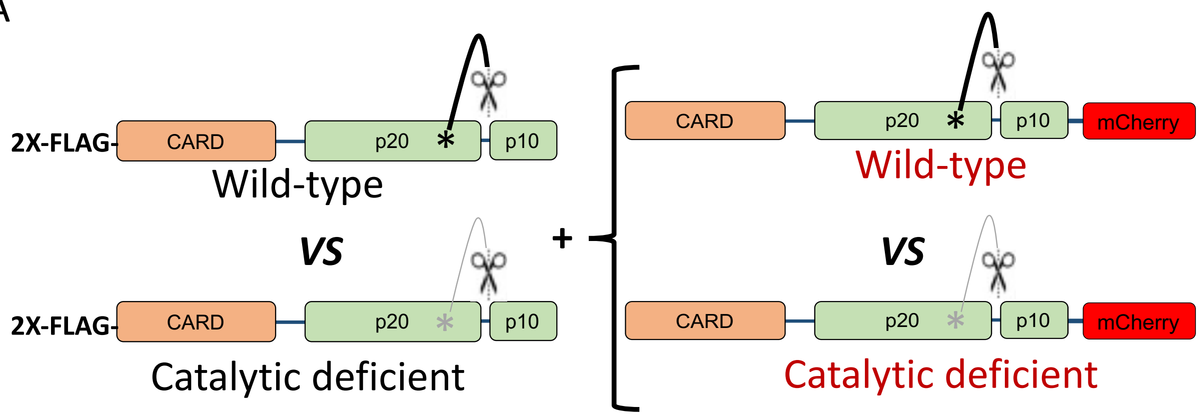


B

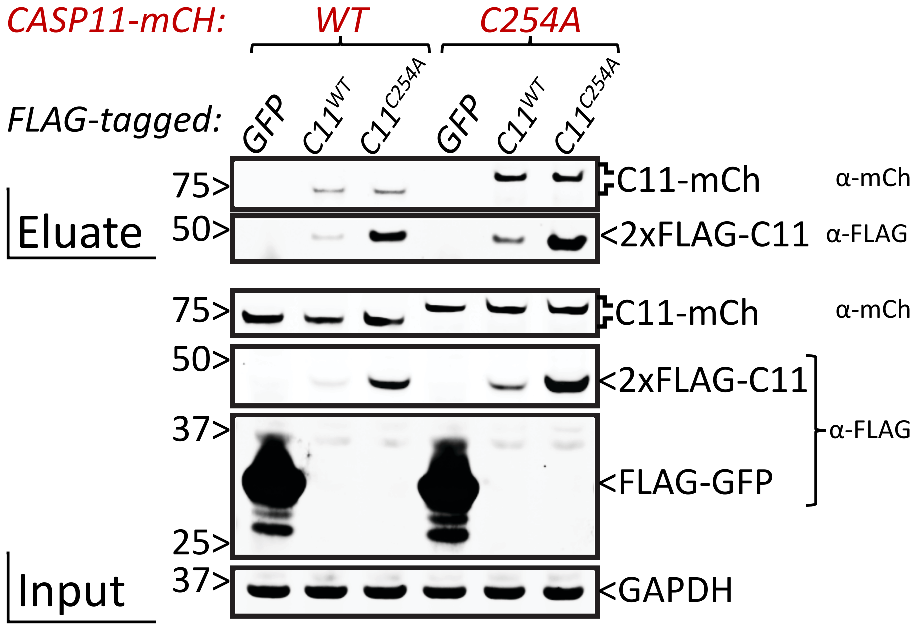

#### Extended Data Figure 4

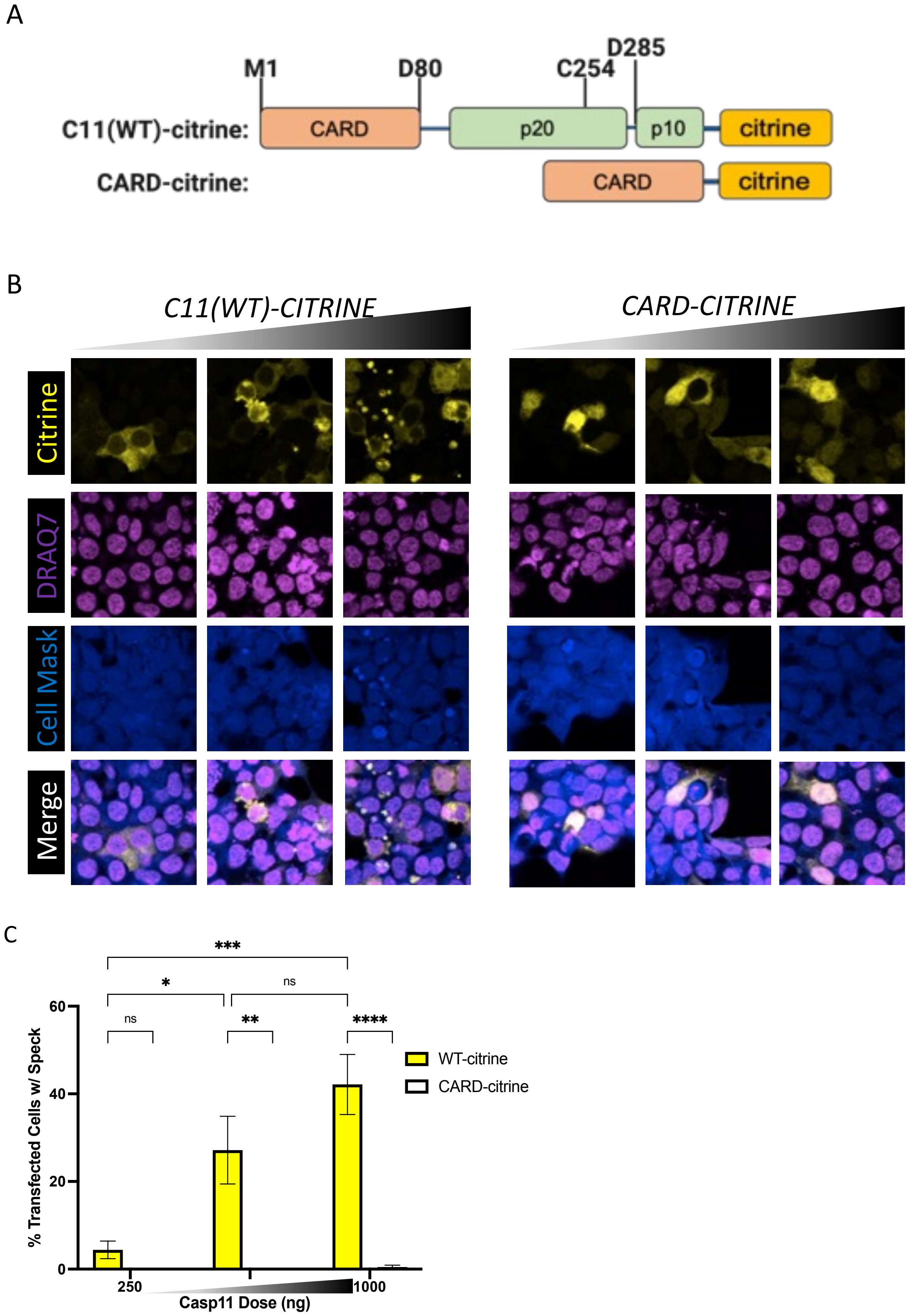
